# Testing the effect of oxytocin on social grooming in bonobos

**DOI:** 10.1101/2021.11.30.468796

**Authors:** James Brooks, Fumihiro Kano, Hanling Yeow, Naruki Morimura, Shinya Yamamoto

## Abstract

Oxytocin has attracted research attention due to its role in promoting social bonding. In bonobos and chimpanzees, the two *Pan* species closely related to humans, urinary oxytocin is known to correlate with key behaviours related to social bonding, such as social grooming in chimpanzees and female-female sexual behaviour in bonobos. However, no study has demonstrated that the administration of oxytocin promotes real-life social interactions in *Pan*, leaving it unclear whether oxytocin is merely correlated with social behaviors or does affect them in these species. To test this, we administered nebulized oxytocin or saline placebo to a group of female bonobos and subsequently observed the change in their gross behavior during free interaction. We found an overall effect of more frequent grooming in the oxytocin condition. However, on the individual level this effect remained significant for only one participant in our follow-up models, suggesting future work should explore inter-individual variation. Our results provide some experimental support for the biobehavioural feedback loop hypothesis, which posits that some functions of the oxytocin system support the formation and maintenance of social bonds through a positive feedback loop; however, further tests with a larger number of individuals are required. Our results, at a minimum, demonstrated that oxytocin affects spontaneous, naturalistic social interactions of at least some female bonobos, adding to accumulating evidence that oxytocin modulates complex social behaviors of *Pan*.

## Introduction

Oxytocin is a hormone neuropeptide conserved through mammalian evolution and plays diverse roles in regulating social behaviors across species. Among non-human great apes, the majority of studies have been conducted through measurement of urinary oxytocin following key social behaviours. Crockford et al. [1] showed that urinary oxytocin levels in wild chimpanzees increase following social grooming, a key socio-positive behavior widely present in nonhuman primates, and proposed that a positive feedback loop through oxytocin may have evolved to support social bonding in this species. Relatedly, Moscovice et al. [2] found that urinary oxytocin levels in wild female bonobos increased following same-sex sexual behaviour, genito-genital (GG) rubbing. Bonobos also increased proximity and coalitionary support among females after GG-rubbing; though it remains unclear if oxytocin played a direct role in these behavioural changes. Other studies have additionally demonstrated that urinary oxytocin in chimpanzees rises after food sharing [3], reconciliation [4], border patrols [5,6], and group hunting [5,7], further suggesting its importance to social bonds and coordination.

In several primate species, studies have demonstrated exogenous oxytocin can impact a wide range of social behaviours (reviewed in [8]). In macaques, several studies have demonstrated that oxytocin alters social gaze, such as increased attention to eyes [9], reduced attention to negative and fearful facial expressions [10] as well as social threats [11], and more gaze following [12]. In one of the first to test the effect of oxytocin in spontaneous social behaviour among multiple macaques, although still confined to primate chairs in a laboratory setting, Jiang and Platt [13] found evidence that oxytocin flattened the dominance hierarchy and enhanced synchrony of mutual gaze. Marmosets similarly showed an increase in attention to eyes [14] following oxytocin administration and an increase in anxiety and vigilance following administration of an oxytocin antagonist [15]. Another study found that oxytocin promoted huddling in marmosets, while an oxytocin antagonist reduced social proximity and huddling [16]. On the other hand, in capuchin monkeys oxytocin was found to reduce food sharing through increasing interindividual distance [17]; the authors interpreted these results as derived from oxytocin’s anxiolytic effect, which increased social distance and thereby decreased opportunities for food sharing [17].

The results of three studies measuring behaviour following oxytocin administration in non-human great apes were mixed. Proctor et al. [18] administered oxytocin to eight chimpanzees individually for one trial each in both saline and oxytocin conditions then observed them in their regular social groups. Although they did not find significant effects for any behaviours measured, the authors note that it may be due to methodological issues, such as in establishing an effective dose of oxytocin for chimpanzees or influence from groupmates who did not receive oxytocin before social interaction. Hall et al. [19] similarly found no effect of oxytocin when chimpanzee dyads were administered oxytocin or saline placebo and subsequently tested in a token exchange task. Each participant chose one of two tokens to exchange and received rewards based on the choice of both participants in distributions based on games such as the prisoner’s dilemma and hawk-dove. However, although this study administered oxytocin to a dyad, the authors reported the same methodological concerns for the oxytocin administration procedure as well as a confound between experimental condition and order. No clear patterns emerged in either the placebo or oxytocin conditions, limiting interpretation of oxytocin’s possible effect. On the contrary to these studies reporting null results, Brooks et al. [20] found that oxytocin enhanced species-typical social gaze, increasing eye contact in bonobos but not chimpanzees, indicating that oxytocin can modulate gaze behaviour. While the species difference in Brooks et al. cannot be attributed to differences in oxytocin administration procedure, it remains unclear whether the lack of effect in Proctor et al. and Hall et al. is due to methodology of oxytocin administration or that exogenous oxytocin fails to significantly affect chimpanzee real-life social interaction.

Therefore, currently there is no study demonstrating that the administration of oxytocin affects spontaneous social interactions of nonhuman great apes, leaving it unclear whether oxytocin does cause any change in key social behaviors of great apes or just is correlated with those behaviors. Moreover, the biobehavioural feedback loop hypothesis suggests that an oxytocin positive feedback loop has evolved to support *Pan* social bonding [1]. Although it is central to this hypothesis that both socio-positive interactions cause oxytocin release and that oxytocin can lead to socio-positive interactions, there is no direct evidence showing that oxytocin promotes any socio-positive interaction in *Pan*.

Given recent progresses in this line of research, it is worthwhile to test whether oxytocin promotes key social behaviours related to social bonding in *Pan* using the updated methods of oxytocin administration. While previous studies with chimpanzees administered oxytocin to one individual or a dyad at a time, and subsequently observed the social interaction between this individual and group mates, we were able to administer oxytocin to whole subgroups of female bonobos simultaneously. For practical reasons, we could only test bonobos (not chimpanzees) in this study design, though similar future work on chimpanzees will also be necessary. In this study, we administered nebulized oxytocin or saline placebo to female bonobos following the methods employed in Brooks et al. [20] and subsequently observed the change in their gross interactive behavior, including grooming and GG-rubbing, as well as other noninteractive behaviours during their free interaction.

## Methods

### Ethics statement

All bonobo participants received regular feedings, daily enrichment, and had ad libitum access to water. No change was made to their daily care routine for the purpose of this study. Apes were never restrained at any point. We carefully considered the safety of the oxytocin administration as in previous studies. Again, we based this decision on the fact that 1) oxytocin is often administered to human children and adults, 2) oxytocin is active for only a short period of time following administration, 3) oxytocin is naturally produced in bonobos and chimpanzees following relevant behaviors [1,2]. and 4) no previous studies administering oxytocin intranasally to chimpanzees or bonobos reported any agonistic interaction [18–20]. All female bonobos were taking birth control (details can be found in supplementary material) and thus no bonobos were pregnant at any time during the course of this experiment. Ethical approval number was WRC-2020-KS014A. This study complied with the American Society of Primatologists Principles for the Ethical Treatment of Non-Human Primates, as well as all applicable laws in the country where it was conducted.

### Participants

Four adult female bonobos at Kumamoto Sanctuary participated in this research. Details about participant ages and rearing histories can be found in supplementary material (Table S1). Animals were not food or water deprived at any time and were given both physical and social environmental enrichment in their daily life. The bonobos live in a dynamic grouping structure where three of the four females are together on any given day, and the fourth is with two male bonobos. These two males were not involved in this study because one male refuses to participate in any oxytocin experiments, and our aim was to test whole groups at a time with the same condition. Three of the females join the male bonobos with varying frequency, while the fourth (Lenore) is always with other female bonobos. Individuals thus had a varying number of trials, with Lenore having the most due to never joining the male group (24 trials), followed by Lolita (20 trials), followed by Louise and Ikela (14 trials each) who are most often with the males. Transfers between groups typically occur in the evening and are kept consistent for at least one day and up to one week. The bonobos therefore had been in the same grouping structure for the whole day prior to the start of experiments.

### Administration procedure

Oxytocin administration procedures followed Brooks et al. [20]. Briefly, oxytocin was dissolved in saline at a concentration of 40IU/mL. The oxytocin solution or placebo control was nebulized into a box using a portable nebulizer (Omron NE-U100) at a minimum rate of 0.25mL/minute, for a cumulative 4 minutes while apes drank juice (thus a total of 40IU or more was nebulized during the administration period). We paused counting the time while apes’ noses were outside the box. Participation to this administration was voluntary. Three of the bonobos could simultaneously participate in oxytocin administration in their typical enclosure, while the fourth (Ikela) preferred to move to another room to participate in the administration procedure. Thus, on days when Ikela was in the group, we first completed the administration procedure with Ikela (accompanied by other participant bonobos), and then returned her to the home enclosure with the other participant bonobos; the other participant bonobos were then administered oxytocin (or saline placebo control). On days when Ikela was not in the group, all participants were administered oxytocin (or saline placebo) in their home enclosure. All group members received the same condition (saline placebo or oxytocin) on any given day of experiments and finished administration procedures within 30 minutes of one another.

One trial was performed in an experimental day. The order of conditions was pseudorandomized such that the same condition (placebo and oxytocin) never occurred more than twice in consecutive trials (experimental days) and that the same number of trials were conducted for placebo and oxytocin condition for each participant and for each grouping structure. We had a minimum of 2 days between trials to avoid any possible carryover effect of oxytocin. On each experimental day, the experiment was performed between 11:00 and 12:15, and the observation window therefore started between 11:30 and 12:45. Experimental days followed the same feeding schedule; bonobos were fed breakfast around 9:00, and additional greenery is available for foraging throughout the day. Experiments took place over five calendar months across two calendar years.

### Observation procedure

Observation began 30 minutes after completion of administration procedures to the last individual, and lasted for one hour. This window was chosen based on previous studies [8], where oxytocin’s effect is typically measured in the window between 30 minutes and 2 hours after completion of administration procedures. In our experiment, the last individual to complete administration procedures was always within 30 minutes of the first individual to finish, and thus all participants were observed for one hour, starting and finishing between 30 minutes and 2 hours following completion of administration procedures on any given day. We chose this window in order to maximize data collection within the active window of oxytocin while ensuring consistency between trials and participants, where observation windows were kept constant at 1 hour per trial and all data points for all individuals were within the 30 minutes to 2 hour window employed in previous studies.

Observation methods combined scan and event sampling. Specifically, every 2 minutes, interindividual proximity was estimated for each dyad into one of four categories; in contact, within arm’s reach (one individual could extend their arm to touch the other), < 3 meters, and > 3 meters. In addition, at the same 2 minute intervals, we coded each individual’s behaviour (grooming - including direction and partner(s), resting, self-directed behaviour, moving, eating). Finally, we recorded all occurrences of play, GG-rubbing, abnormal behaviour (in this group primarily regurgitation and reingestion), and aggression towards groupmates (including displays).

We additionally recorded any agonistic or socio-sexual behaviour during the administration procedures and during the 30 minutes before the formal observation window to check if the presence of such interactions could account for our results. No agonistic behaviour was observed, and GG-rubbing occurred on 4 trials during administration procedure, on 2 trials before oxytocin condition and 2 trials before saline placebo condition.

### Analysis

All analyses were conducted in R [27]. Behavioural scan data was analyzed with binomial GLMMs (Generalized Linear Mixed Models) with package lme4 [28], where each individual at each sampling point was characterized as either engaged in (1) or not engaged in (0) a given behaviour. The model included a fixed effect of condition as a test effect and also fixed effects of time, the square of time (time^2^) and grouping structure (where a unique value was given for each possible combination of individuals) as control effects. The model also included random effects of participant and day (a factor with a unique value for each experimental day), as well as random slopes of each fixed effect for each random effect. Numeric effects were z-transformed to have a mean of 0 and the standard deviation of 1. Random slope structure was kept maximal, except that the interaction between random slopes and intercepts was removed due to issues with convergence [29]. The model structure was thus: behaviour ~ condition + time + time^2^ + group + (1 + group + condition + time + time^2^1 | individual) + (1 + group + condition + time + time^2^1 |day).

For all GLMMs we checked model stability by comparing our models to those which excluded levels of the random effects one at a time. We additionally calculated odds ratio (OR) estimates and 95% confidence intervals for all significant effects. For the grooming data, we analyzed rates of active grooming (giving or mutual grooming) on the individual level, where receiving grooming was valued as 0 for not actively grooming another individual. Given considerable inter-individual variation observed by visual inspection of the data, we additionally tested each individual with the same model structure for significant effects (without the random effect of participant). We here used a significance threshold of 0.0125 (0.05/4) to correct for multiple testing at the individual level. Finally, we ran models excluding each level of participant to check the overall model stability.

Proximity data was analyzed using a CLMM (cumulative link mixed model) on ordinal data using the package “ordinal” [30]. Fixed and random effect structures were the same as those in the behavioural scan data analysis, except for the individual participant variable was replaced by dyad (a unique value for each dyad), and the addition of two random effects to represent the two individuals within a dyad (randomly distributed as individual variable 1 and 2).

All occurrence data was analyzed with a binomial GLMM, where each individual for each day was characterized as having engaged in (1) or not engaged in (0) a given behaviour. The fixed and random effect structure was the same as in the scan behaviour models, except for the time variables were removed due to data being summarized across a given observation day. Model syntax for all model types can be found in supplemental material.

For all models, statistical significance of effects was calculated using a likelihood ratio test (using the “drop1” function in R).

## Results

Our model with active grooming as response revealed a significant effect of condition (oxytocin, saline placebo; *β* = 1.16, SE = 0.49, χ^2^ = 5.47, *p* = 0.019, OR = 3.18 (95% CI: 1.23, 8.23); Table 1, Figure 1). In our stability check analysis excluding each participant one-by-one in the model, the effect of oxytocin on grooming was significant in all models except for that excluding Louise (excluding Ikela: *β* = 0.79, SE = 0.31, χ^2^ = 5.27, *p* = 0.022; excluding Lenore: *β* = 1.28, SE = 0.64, χ^2^ = 8.97, *p* = 0.0027; excluding Lolita: *β* = 0.88, SE = 0.36, χ^2^= 5.06, *p* = 0.025; excluding Louise: *β* = 0.59, SE = 0.37, χ^2^= 2.34, *p* = 0.13). In our individual-level analysis of grooming, we found a significant effect of condition only for Louise using a significance threshold of 0.0125 (Ikela: *β* = 0.86, SE = 1.51, χ^2^ = 0.24, *p* = 0.63; Lenore: *β* = 0.79, SE = 0.35, χ^2^ = 4.61, *p* = 0.032; Lolita: model could not run (as Lolita never actively groomed in our dataset); Louise: *β* = 1.15, SE = 0.40, χ^2^ = 9.09, p = 0.0026).

**Figure 1:**
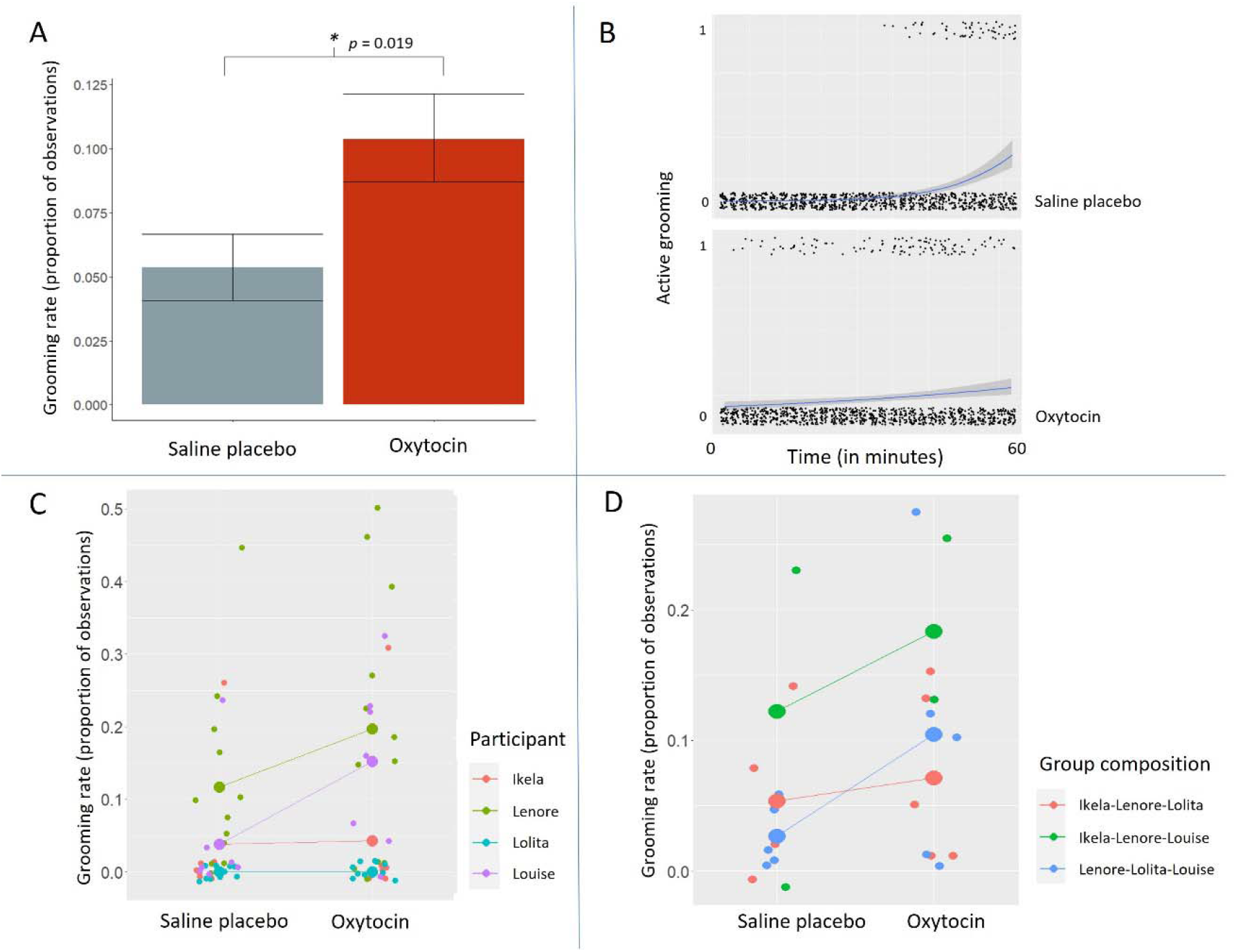
Rates of active (giving) grooming in the oxytocin and saline placebo conditions over the observation window. A) Absolute rates of grooming across all trials and participants B) Time course of grooming by condition, 1 represents giving grooming and 0 represents not giving grooming C) Grooming rates by participant and condition (small circles represent grooming rates by trial, large circles represent mean grooming rates across trials) D) Grooming rates by group structure by condition (small circles represent grooming rates by trial, large circles represent mean grooming rates across trials).

**Table 1;.** Details of grooming and self-directed behaviour models with significant terms in bold.

| Response | Term | Estimate±SE | $\chi^2$ | df | p |
| --- | --- | --- | --- | --- | --- |
| Active grooming | (Intercept) | -15.57±6.29 | | | |
|  | Test predictors: |  |  |  |  |
|  | **Condition** | **1.156±0.49** | **5.47** | **1** | **0.019\*** |
|  | Control predictors: |  |  |  |  |
|  | *Group* |  | 4.73 | 2 | 0.094. |
|  | *Ikela-Lenore-Louise* | *12.61±6.29* |  |  |  |
|  | *Lenore-Lolita-Louise* | *8.14±6.70* |  |  |  |
|  | **Time (time + time^2)** | **1.59±0.44** | **9.15** | **1** | **0.0025\*** |
| Self-directed behaviour | (Intercept) | -2.81±0.60 |  |  |  |
|  | Test predictors: |  |  |  |  |
|  | **Condition** | **-0.33±0.14** | **5.16** | **1** | **0.018\*** |
|  | Control predictors: |  |  |  |  |
|  | Group |  | 3.09 | 2 | 0.21 |
|  | Ikela/Lenore/Louise | -0.094±0.53 |  |  |  |
|  | Lenore/Lolita/Louise | 0.75±0.53 |  |  |  |
|  | **Time** | **-0.31±0.13** | **4.42** | **1** | **0.035\*** |

There was also a significant reduction in self-directed behaviour in the oxytocin compared to placebo condition (*β* = −0.33, SE = 0.14, χ^2^= 5.16, *p* = 0.018, OR = 0.72 (95% CI: 0.54, 0.94); Table 1). In our stability check analysis excluding each participant one-by-one in the model, the effect of oxytocin on self-directed behaviour was significant in the model excluding Lenore and the model excluding Lolita (excluding Ikela: *β* = −0.26, SE = 0.13, χ^2^ = 3.52, *p* = 0.061; excluding Lenore: *β* = −0.41, SE = 0.14, χ^2^= 8.89, *p* = 0.0029; excluding Lolita: *β* = −0.51, SE = 0.16, χ^2^ = 9.81, *p* = 0.0017; excluding Louise: *β* = −0.34, SE = 0.17, χ^2^ = 3.65, *p* = 0.056). In our individual-level analysis, we found a significant effect only for Ikela using a significance threshold of 0.0125 (Ikela: *β* = −0.71, SE = 0.21, χ^2^ = 12.19, *p* = 0.00048; Lenore: *β* = −0.15, SE = 0.28, χ^2^= 0.27, *p* = 0.60; Lolita: *β* = −0.59, SE = 0.24, χ^2^ = 5.58, *p* = 0.018; Louise: *β* = −0.58, SE = 0.41, χ^2^ = 2.48, *p* = 0.11).

There were no significant differences in interindividual proximity (*β* = −0.15, χ^2^= 0.32, *p* = 0.57), frequency of the abnormal behaviour regurgitation and reingestion (*β* = −1.05, SE = 0.66, χ^2^ = 2.65, *p* = 0.10), or rate of rest (*β* = −0.10, SE = 0.18, χ^2^= 0.34, *p* =0.56) between the oxytocin and placebo condition. Bonobos engaged in GG-rubbing only once (oxytocin condition) and displayed no aggression toward groupmates or any bouts of play during the observation period. See supplementary material Table S2 for full details of all models.

## Discussion

We found that the grooming rates of captive female bonobos were higher in the oxytocin compared to saline placebo condition, consistent with the predictions of the biobehavioural feedback loop hypothesis of oxytocin in bonobo social bonding. However, when conducting individual-level analyses this was only significant after correcting for multiple testing in one participant (Louise). There was no significant effect of oxytocin on inter-individual proximity, suggesting the increased rate of grooming is not merely a consequence of increased proximity. The bonobos also engaged in self-directed behaviour less in the oxytocin compared to placebo condition (though again this was significant on an individual level in just one participant - Ikela), which is potentially related to its anxiolytic effect, though it should be noted we did not distinguish between kinds of self-directed behaviours such as self-scratching or self-grooming which may have different relations to stress. The proportion of rest and frequency of regurgitation and reingestion did not differ between conditions, while GG-rubbing, play, and aggression were rarely or never observed during our 1-hour observation window, likely due to low overall tension, precluding formal analysis.

Despite our overall model showing an increase in grooming in the oxytocin condition, our individual-level analyses revealed only one individual, Louise, a 48-year-old female bonobo relatively dominant to other groupmates. Lenore, a 38-year-old, also groomed groupmates more in the oxytocin compared to the saline condition, but was not significant after correcting for multiple testing. Ikela, a 29-year-old, groomed slightly more, but this effect was also not significant. Lolita was never observed actively grooming throughout our experiment, and thus her rate of grooming was completely unchanged by oxytocin. Future studies are needed to examine what factors drive these potential individual differences. It is also important to note that given social grooming necessarily requires a grooming partner, and given our experimental design administering the same condition to all group members simultaneously, oxytocin’s effect on the group may not be entirely reducible to the individual level. Oxytocin may as a whole promote a group dynamic more conducive to grooming, which is measurable in the behaviour of certain key individuals. This possibility can be directly tested by administering oxytocin compared to saline placebo only to Louise, and always saline placebo to others, and examining if the same effect is found.

Although we addressed some previous methodological issues, there are several important limitations in this study. Due to limited possibility of testing, enclosures suitable for detailed observation, and some apes’ willingness to join experiments, the sample was limited to four adult female bonobos. Moreover, previous work has indicated sex-specific effects of oxytocin [31–33], and thus it remains unclear whether our results can be generalized to different sex pairs, though it should be noted that Crockford et al. [1] did not find significant differences between female-female, female-male, and male-male dyads in urinary oxytocin level following grooming in wild chimpanzees. We also could not investigate the possible effect of different dominance rank, rearing history, age, or genetic background contributing to differences in the amount of change between conditions across individuals, which should be directly explored in the future. Moreover, the small number of participants did not enable us to test the effect of existing social bond strength and relatedness among groupmates, which may interact with the observed increase in grooming.

Finally, while we found an effect of oxytocin on rates of social grooming at least in some individuals, we did not find any effect on rates of GG-rubbing. An increase in urinary oxytocin following GG-rubbing was reported by Moscovice et al.’s [2]. We observed GG-rubbing just once in the experiment’s observation window. GG-rubbing is typically infrequent in our study group of bonobos, particularly under normal conditions in their home environment as in our main observational window. Floor effects may thus be responsible for the lack of an effect, and future studies focused on contexts where GG-rubbing is more likely to occur (e.g., feeding, reunions) will be necessary to determine whether or not oxytocin increases propensity to engage in GG-rubbing in female bonobos.

In conclusion, we found that exogenous oxytocin promotes social grooming in at least some female bonobos when administered to the whole group. Although much future work is necessary, our results demonstrate that oxytocin does affect a socio-positive behavior of a *Pan* species during spontaneous, naturalistic social interactions, filling some gap between both previous field studies on ape urinary oxytocin and experimental administration studies on non-ape species. Moreover, although limited, our finding offers some experimental evidence for the biobehavioural feedback loop hypothesis for oxytocin in bonobo social bonding when combined with previous research showing increased urinary oxytocin in bonobos and chimpanzees following socio-positive interaction [1,2] (note also that while peripheral measures such as urinary measurement allow non-invasive data on primates, central oxytocin release following specific social behaviours has also been found in rodents [34]). Future work should test a larger number of individuals to test potential differences in oxytocin’s effect between species, should examine inter-individual variation with respect to social closeness and centrality, and should study how social contexts such as feeding tension interact with this effect. Our results, at minimum, demonstrate that oxytocin can affect socio-positive behaviour in at least some bonobo individuals during natural social interactions, adding to accumulating evidence on its importance to *Pan* sociality.

## Acknowledgements

We thank the bonobos at Kumamoto Sanctuary for participating in this study. We also thank Etsuko Nogami and the other caretakers at Kumamoto Sanctuary for their support throughout this project, Dr. Satoshi Hirata for his advice and support, and Dr. Yutaro Sato for helping in data collection. This study was funded by Japan Society for the Promotion of Science (KAKENHI #21J21123 to J.B., #19H01772 and #20H05000 to F.K., and #19H00629, #22H05653, and #22H04451 to S.Y.)

## Notes

### Competing Interest Statement

The authors have declared no competing interest.

### Summary of Updates

Introduction and discussion revised to clarify and simplify main points. Added some additional information.

