## supplementary material for "Testing the effect of oxytocin on social grooming in bonobos"

Table S1. Participant information. In the rearing history column, ‘Mother’ indicates the individuals reared by their biological mothers, and ‘Nursery-peer’ indicates the individuals reared by human caregivers and conspecific peers.

| Name | Sex | Age | Rearing history | Days (hours) observed |
| --- | --- | --- | --- | --- |
| Ikela | F | 29 | Nursery-peer | 14 |
| Lenore | F | 38 | Mother | 24 |
| Louise | F | 48 | Nursery-peer | 14 |
| Lolita | F | 31 | Nursery-peer | 20 |

Birth control method:

Lenore: Ovariectomy
Lolita & Ikela: Norethisterone 1.0mg + ethinylestradiol 0.035mg once a day, with a schedule of 3-weeks on and 1-week off
Louise: Norethisterone 1.0mg every day

Model syntax:

Behavioural scan data:

Rate or behaviour ~ condition + group + time_+ time^2

+ (1 + condition + group + time + time^2||individual)

+ (1 + condition + group + time + time^2||day)

Proximity data:

Proximity category ~ condition + time + time^2 + group

+ (1 + condition + group + time + time^2 ||dyad)

+ (1 + condition + group + time + time^2||day)

+ (1 + condition + group + time + time^2||ID1)

+ (1 + condition + group + time + time^2||ID2)

*note that the “ordinal” R package does not support the || shortcut to remove the interactions between random slopes and random intercepts and was done manually

All occurrence data:

Rate of behaviour ~ condition + group

+ (1 + condition + group||individual)

+ (1 + condition + group||day)

Table S2: Detailed model results

| Response | Term | Estimate ± SE | *χ^2^* | *df* | *p* |
| --- | --- | --- | --- | --- | --- |
| Grooming | (Intercept) | -15.57±6.29 |  |  |  |
|  | Test predictors: |  |  |  |  |
|  | **Condition** | **1.156±0.49** | **5.47** | **1** | **0.019*** |
|  | Control predictors: |  |  |  |  |
|  | *Group* |  | *4.73* | *2* | *0.094.* |
|  | *Ikela/Lenore/Louise* | *12.61±6.29* |  |  |  |
|  | *Lenore/Lolita/Louise* | *8.14±6.70* |  |  |  |
|  | **Time** | **1.59±0.44** | **9.15** | **1** | **0.0025*** |
| Self-directed behaviour | (Intercept) | -2.81±0.60 |  |  |  |
|  | Test predictors: |  |  |  |  |
|  | **Condition** | **-0.33±0.14** | **5.16** | **1** | **0.018** |
|  | Control predictors: |  |  |  |  |
|  | Group |  | 3.09 | 2 | 0.21 |
|  | Ikela/Lenore/Louise | -0.094±0.53 |  |  |  |
|  | Lenore/Lolita/Louise | 0.75±0.53 |  |  |  |
|  | **Time** | **-0.31±0.13** | **4.42** | **1** | **0.035** |
| Resting | (Intercept) | 0.86±0.43 |  |  |  |
|  | Test predictors: |  |  |  |  |
|  | Condition | -0.10±0.18 | 0.34 | 1 | 0.56 |
|  | Control predictors: |  |  |  |  |
|  | Group |  | 3.76 | 2 | 0.15 |
|  | Ikela/Lenore/Louise | -0.36±0.56 |  |  |  |
|  | Lenore/Lolita/Louise | -0.81±0.40 |  |  |  |
|  | Time | -0.20±0.23 | 0.72 | 1 | 0.40 |
| Regurgitation/Reingestion | (Intercept) | -2.39±1.29 |  |  |  |
|  | Test predictors: |  |  |  |  |
|  | Condition | -1.05±0.66 | 2.65 | 1 | 0.10 |
|  | Control predictors: |  |  |  |  |
|  | Group |  | 2.47 | 2 | 0.29 |
|  | Ikela/Lenore/Louise | -19.01±4854 |  |  |  |
|  | Lenore/Lolita/Louise | 0.45±1.58 |  |  |  |
| Proximity (using CLMM) | Test predictors: |  |  |  |  |
|  | Condition | -0.15 | 0.32 | 1 | 0.57 |
|  | Control predictors: |  |  |  |  |
|  | Group |  | 2.06 | 2 | 0.36 |
|  | Ikela/Lenore/Louise | -0.71 |  |  |  |
|  | Lenore/Lolita/Louise | -0.39 |  |  |  |
|  | **Time** | **-4.23** | **5.05** |  | **0.025*** |
